# Infection avoidance behaviour: female fruit flies adjust foraging effort in response to internal and external cues of viral infection

**DOI:** 10.1101/039750

**Authors:** Pedro F. Vale, Michael D. Jardine

## Abstract

Infection avoidance behaviours are the first line of defence against pathogenic encounters. Behavioural plasticity in response to internal or external cues can therefore generate potentially significant heterogeneity in infection. We tested whether *Drosophila melanogaster* exhibits infection avoidance behaviour during foraging, and whether this behaviour is modified by prior exposure to Drosophila C Virus (DCV) and by the risk of DCV encounter. We examined two measures of infection avoidance: (1) the motivation to feed in the presence of an infection risk and (2) the preference to feed on a clean food source over a potentially infectious source. While we found no clear evidence for preference of clean food sources over potentially infectious ones, female flies were less motivated to feed when presented with a risk of encountering DCV, but only if they had been previously exposed to the virus. We discuss the relevance of behavioural plasticity during foraging for host fitness and pathogen spread.

## Introduction

Hosts vary considerably in their ability to acquire and transmit infection ^1^^-^^3^, and much of this variation is caused by differences in the contact rate between susceptible individuals and sources of infection^4,5^. For example, viruses of *Drosophila* fruit flies are not only widely distributed, they also show very broad host range^6^. Given the high viral prevalence of pathogens in natural environments, mounting a timely and efficient immune response to all possible pathogenic challenges would be physiologically costly and ultimately ineffective. Hosts capable of reducing the probability of contacting parasites, infected conspecifics or infectious environments can therefore not only prevent the deleterious effects of infection, but also circumvent the undesirable energetic costs of immune responses, including immunopathology ^4,7^. Avoiding infection is therefore the first line of non-immunological defence against infection^8^, and is known to occur across a broad range of host taxa, including humans ^7,9^.

Like most traits, infection avoidance behaviours are likely to vary according to the context of infection, and pathogens are major drivers this context ^4,7,9-11^ Pathogens may alter host responses in two ways. By altering the immunophysiology of the host during infection, pathogens can alter host behaviour^12,13^. Pathogens also modify the host external environment by increasing the likelihood of exposure to novel infections, and these external cues of infection risk are also known to influence host behavioural responses ^4,7^. Understanding variation in infection avoidance behaviours therefore provides an important functional link between the neurological, behavioural and immunological processes that together govern the spread of disease ^12^.

Insects are ideal systems to investigate the interplay between infection and behaviour ^12,14^. The fruit fly *Drosophila* is especially amenable to these studies, as it is one of the best developed model systems for host-pathogen interactions ^15^ and behavioural ecology and genetics ^16,17^. One of the most studied pathogenic interactions in *Drosophila* is the host response to systemic and enteric infection with Drosophila C Virus (DCV)^18,19^. DCV is a horizontally transmitted +ssRNA virus that naturally infects the fly gut ^19^^-^^21^, causing intestinal obstruction, severe metabolic dysfunction and eventually death ^22,23^. As a consequence of its pathology, female flies infected with DCV are also known to exhibit behavioural modifications, such as reduced locomotion and increased sleep ^24^. The Drosophila-DCV interaction therefore offers a powerful system to investigate the ecological consequences that may arise from the physiological and behavioural effects of enteric viral infections.

In the present study we used a combination of controlled experimental infections and foraging choice assays, to test whether adult *D. melanogaster* are able to avoid potentially infectious environments when foraging for food, and if avoidance behaviour is modified in response to virus exposure history and to different risks of acquiring DCV infection. We find evidence for avoidance behaviours in the form of reduced motivation to feed according to the risk of infection. However, these effects were only present in female flies previously exposed to DCV, indicating potentially important sexual dimorphism in infection avoidance.

## Materials and methods

### Fly and virus stocks

All flies used were from a long-term laboratory stock of Wolbachia-free *Drosophila melanogaster* Oregon R line, maintained on Lewis medium in standard conditions: 25°C, with a 12:12h light:dark cycle. Fly stocks were routinely kept on a 14-day cycle with non-overlapping generations under low larval densities. The DCV culture used in this experiment was grown in Schneider Drosophila Line 2 (DL2) as described in ^24^. Ten-fold serial dilutions of this culture (diluted in Ringers buffer solution) were aliquoted and frozen at −80°C for long-term storage before use.

### Virus exposure

Flies used in the foraging choice assays were obtained by preparing 10 vials of Lewis medium and yeast containing ten mated females. Flies were allowed to lay eggs for 48 hours resulting in age-matched progeny reared in similar larval densities. To test the effect of previous exposure to virus on avoidance behaviour during foraging, We exposed these progeny to DCV via the oral route of infection two to three days after eclosion. Oral DCV infection causes small but significant reduction in fly survival^19^ and in we have found that orally infected flies experience changes affects fly mortality, fecundity, fecal shedding (Vale, unpublished data), activity and sleep^24^. Briefly, single-sex groups of 20 flies were placed in vials containing agar previously sprayed with DCV (“exposed” to 50 µl of 10^8^ viral copies/ml) or the equivalent volume of Ringers buffer solution as a control (“not exposed”). This procedure produced 10 replicate vials of either healthy or virus-exposed male or female flies. The viral dose used here was lower than previously reported methods^19^, so we first tested this dose was sufficient to result in viable DCV infections by measuring changes in virus titres and fly survival in separate experiments (Fig. 1). Fly survival was monitored on 5 replicate groups of 10 OreR female flies per vial for 11 days following oral exposure. To measure changes in DCV titre, twenty-five, 2-3 day-old female flies were individually housed in vials previously sprayed with DCV as described above for 3 days. Five flies were collected 1, 3, 6, 9 or 13 days after exposure and total RNA was extracted from flies homogenised in Tri Reagent (Ambion), reverse-transcribed with M-MLV reverse transcriptase (Promega) and random hexamer primers, and then diluted 1:10 with nuclease free water. qRT-PCR was performed on an Applied Biosystems StepOnePlus system using Fast SYBR Green Master Mix (Applied Biosystems). We measured the relative fold change in DCV RNA relative to *rp49*, an internal *Drosophila* control gene, calculated as 2^-ΔΔCt^as described in ^25^.

**Fig. 1.**
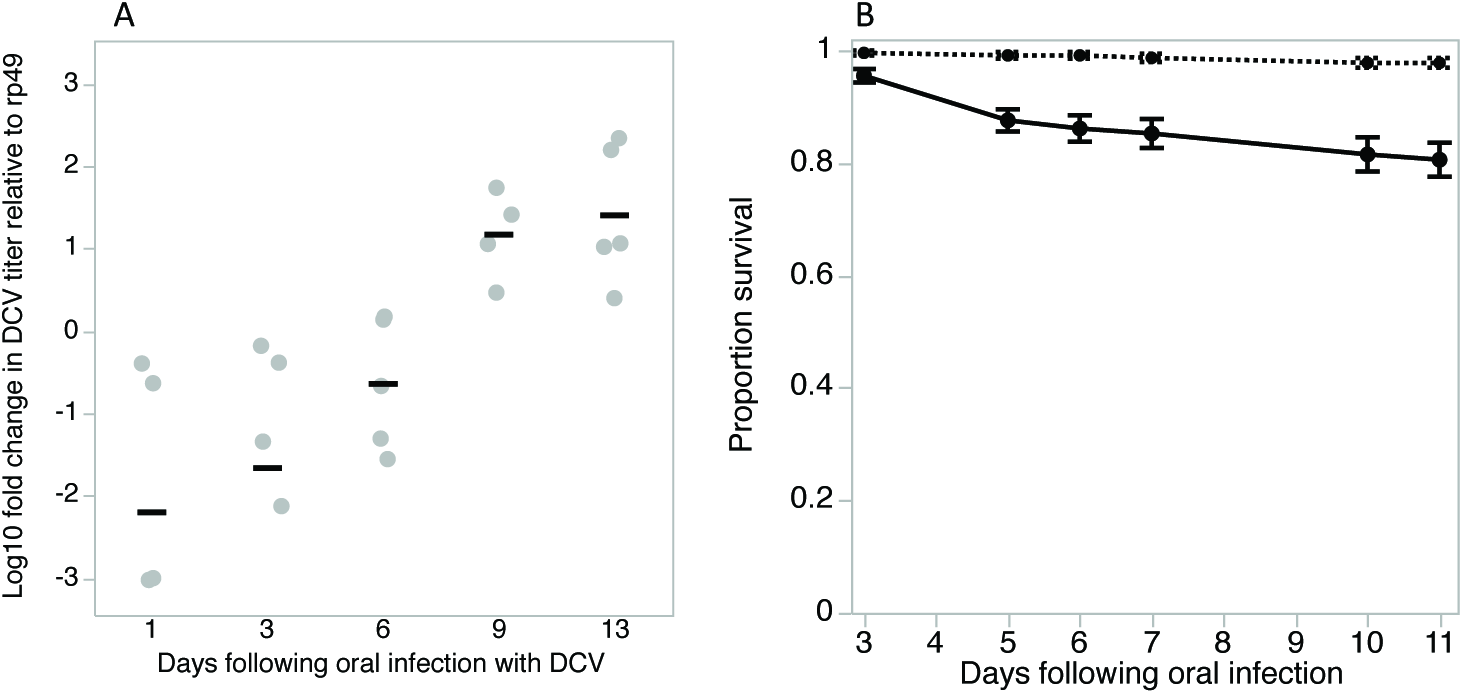
Exposing flies to DCV by placing them in DCV-contaminated vials for three days resulted in flies acquiring replicating virus as shown by the increase in DCV titres over time (Fig. 1A). Grey points shown are individual replicate female flies, black bars are mean titres. This orally acquired DCV infection had a moderate effect on fly survival (full black line) compared to uninfected control flies (dotted line) (Fig. 1B).

### Foraging choice assays

Following 3 days of virus exposure, we setup independent foraging choice assays in cages - cylindrical transparent plastic containers (12 cm in diameter) containing two equally spaced plastic vials of standard Lewis fly medium supplemented with dry yeast. For each combination of “DCV exposed” and “not exposed” male or female flies, we set up two sets of cages to simulate different risks of infection: a “no risk” environment, with two clean vials (sprayed with sterile Ringers solution), and a “high-risk” environment where one of the vials was sprayed with DCV, as described above. Six replicate 20-fly groups were allocated to the “high-risk” chambers and four replicates to the “no risk” chambers, resulting in a total of 40 independent foraging choice cages. Flies were added to the chamber from a neutrally placed hole in the lid, and the number of flies that settled on each vial was recorded every 30 minutes for five hours. Care was taken to randomise the position of the cages so that the orientation of the light did not influence the choice of the flies in any systematic way.

### Statistical Analysis

To measure infection avoidance, we took two approaches. First, we hypothesised that the motivation to feed would be lower in environments where the risk of infection is higher^7^. We therefore compared the motivation to feed between the “no risk” and “high-risk” cages, measured as the proportion of flies inside each replicate cage that chose to feed on any of the provided food sources. We also asked whether flies that chose to feed showed any evidence of avoiding potentially infectious food sources. For this analysis we focussed on the “high risk” cages and recorded the proportion of flies choosing the clean food source over the infectious food source in each replicate cage. In both analyses of motivation to feed and infection avoidance, data on the proportion of flies choosing each food source within each replicate cage were analysed with a generalised linear model assuming binomial error and logit link function, and included fly sex, previous exposure and infection risk as fixed effects. Replicate cage was included as a random effect, nested within treatments. We also analysed the average motivation to feed and infection avoidance across all time points, in a model that inlcuded “time” as a random effect. Treatment specific contrasts were used to test the significance of pairwise comparisons. Analyses were carried out using JMP 12 ^26^.

## Results and Discussion

The ability to detect and discriminate between clean and potentially infectious environments is vital to avoid the adverse consequences of infection. In this study we tested if infection avoidance behaviour in *Drosophila melanogaster* is modified by its previous exposure to a viral pathogen and by the risk of infection with that same pathogen when encountered during foraging. Viral exposure prior to the behavioural assay was achieved by placing flies in a DCV contaminated environment for 3 days, allowing flies to acquire DCV infection orally. DCV acquired through the oral route using this protocol continued to replicate within the fly, increasing by 10-100 fold by day 13 following oral exposure (F_4,19_ = 8.78, p=0.0003; Fig. 1A) and resulted in up to 20% mortality within this period (Fig. 1B).

In the foraging choice assay, only a fraction of flies chose either of the food sources provided, and this motivation to feed increased over time for flies in all treatment groups (χ^2_1_^= 11.00, p=0.001; Fig. 2A). The rate at which motivation increased differed between sexes (Time × Sex interaction, χ^2_1_^= 12.47, p=0.0004), and on average female flies showed greater motivation to feed than males (χ^2_1_^= 5.01, p=0.025), with 67% of female and 36% of male flies making a choice to feed on any of the provided substrates during the observation period. Once flies had made the choice to feed on one of the provided food sources, the choice between a clean and a potentially infectious food source was not affected by previous exposure to DCV (previous exposure, χ^2_1_^= 0.513, p=0.47) in either male or female flies (sex, χ^2_1_^= 0.595, p=0.44).

**Fig. 2.**
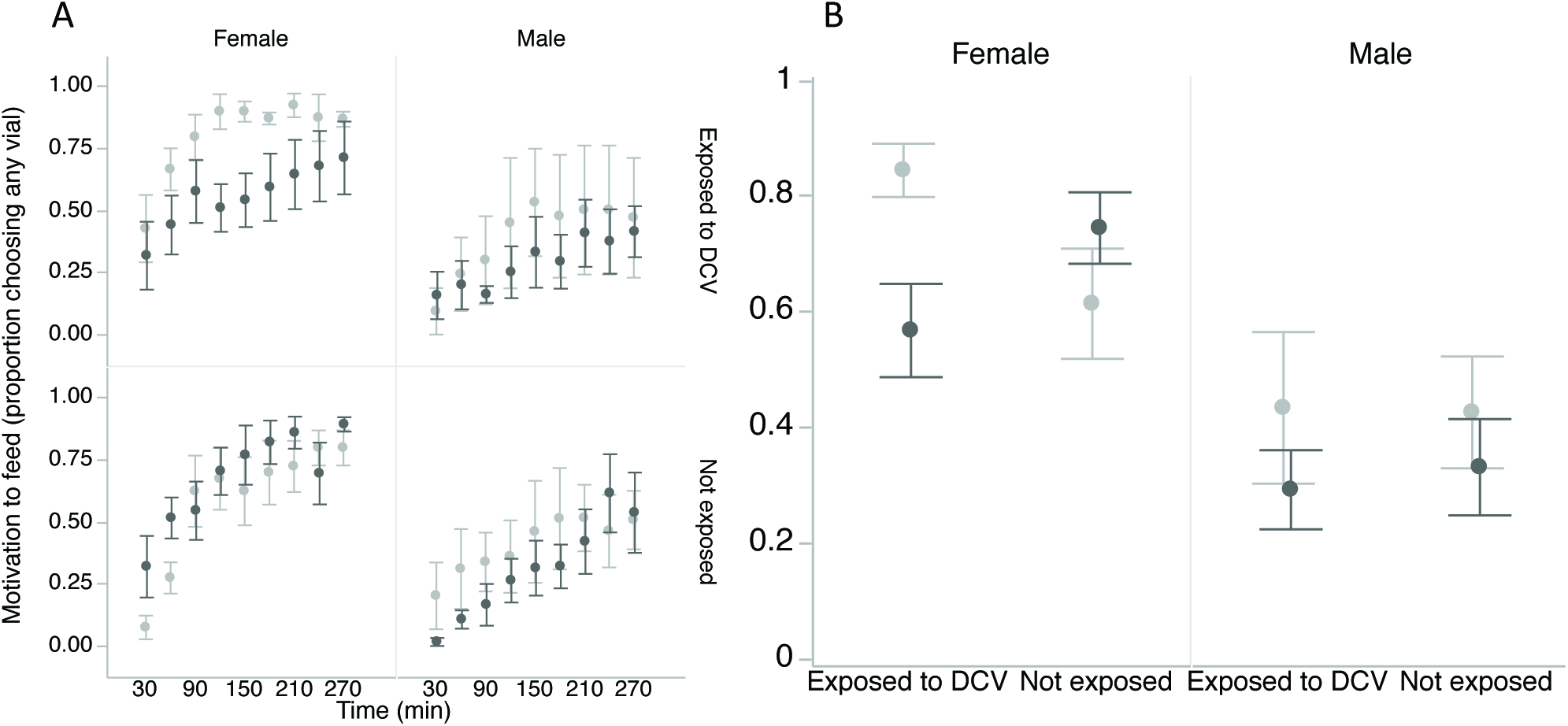
Single sex-groups of flies that had been previously exposed either to DCV or to a sterile inoculum were tested in a 'no-risk' environment (choice between two clean vials; light grey) or a high-risk environment (choice between a clean vial and a DCV-contaminated vial; dark grey). The motivation to feed, measured as the proportion of flies in the cage that fed on any of the provided food sources, increased over time (Fig 2A). Fig 2B shows the average of motivation to feed taken across the whole observation period for each combination of fly sex, prior DCV exposure and current exposure risk (no-risk environment (light grey) or a high-risk environment (dark grey). Data show means ± SE.

Across the entire observation period, the motivation to feed differed between sexes, and depended both on their previous exposure and on their current risk of infection (Sex × risk of infection × Previous exposure interaction, χ^2_1_^= 21.82, p<0.0001). The proportion of males choosing any food substrate did not vary with previous exposure to DCV in either high-risk (χ^2_1_^= 2.21, p=0.137) or no-risk environments (χ^2_1_^= 0.09, p=0.764; Fig. 1). In female flies however, previous exposure and current infection risk affected the motivation to feed on the provided food sources. When there was no risk of infection (Fig. 2B, light grey bars) the motivation to feed was greater in females that were previously exposed to DCV than in otherwise healthy, non-exposed females (χ^2_1_^= 104.11, p<0.001). Among females that were previously exposed to infection, we found that the presence of a risk of acquiring infection resulted in lower foraging effort - with just over 50% of flies making the choice to feed - compared to females in cages where there was no risk of acquiring infection, where over 80% of flies made the choice to feed (Fig. 2B; χ^2_1_^= 168.48, p<0.001).

In addition to responding to external cues of infection (infection risk), internal physiological cues (in this case, previous exposure to DCV) may also modify avoidance behaviour. Behavioural modifications due to infection are widely reported among animals ^9,27^, and can be classified into (i) parasitic manipulation that enhances parasite transmission^9^ (ii) sickness behaviours that benefit the host by conserving energetic resources during infection ^13^, or (iii) side-effects of pathogenicity that do not benefit the host or the parasite ^27^. Female flies infected orally with DCV are known to experience increased lethargy and sleep ^24^, so these effects could also explain the reduced feeding activity we detected in female flies that had been previously exposed to DCV. Another potential explanation for reduced motivation to feed in previously exposed flies is infection-induced anorexia^28^, a commonly described sickness behaviour^13^. However, it is unlikely that a lower motivation feed is simply a symptom of a “sick” fly because it varied according to the risk of infection, and even reached 80% in exposed flies when foraging in a no risk environment (Fig. 2). This suggests that flies are actively avoiding contact with the potentially contagious food source by lowering their foraging effort.

The higher motivation to feed of some female flies when the risk of infection was absent (Fig. 2) suggests flies were able to identify external cues of infection risk. Identifying infection cues is a general prerequisite of avoidance behaviours and occurs across a wide range of different taxa. For example, lobsters are known to detect and avoid virus-infected conspecifics ^29^; fruit flies and nematodes are capable of avoiding pathogenic bacteria^30,31^; gypsy moth larvae are able to detect and avoid virus-contaminated foliage ^14^; sheep have been found to prefer to graze in parasite-poor patches ^32^; and it is has been argued that the disgust response in humans has evolved because it decreases contact with potential infection ^33^. It is unclear how flies are able to identify food sources contaminated with a viral pathogen. In *Drosophila* and *C. elegans* avoidance of pathogenic bacteria is enabled by evolutionary conserved olfactory and chemosensory pathways ^30,31,^ while avoidance of parasitic wasps appears to be mainly enabled by the visual sensory system ^34^. While avoiding virus infected conspecifics is probably driven by visual cues of infection ^29^, it remains unclear how virus-contaminated environments may trigger a lower motivation to feed in *Drosophila.*

The fact that only female flies demonstrated avoidance is an indication that any potentially adaptive effects of avoiding infection may be related to oviposition, which coincides with feeding. For flies previously exposed to DCV, avoiding infection would not confer substantial direct benefits given the physiological and behavioural costs of this infection ^22^^-^^24^, but would however reduce the exposure of future offspring to infection. While flies previously exposed to DCV do not appear to immune primed following an initial viral exposure ^35^, our results point to a sort of behavioural priming, where females previously exposed to infection avoid foraging in potentially infectious environments.

In summary, using a combination of experimental infections and behavioural assays, we find evidence for infection avoidance in *Drosophila* in the form of reduced motivation to feed, which was most pronounced when flies were faced with an increased risk of encountering an infectious food source. However, these effects were only present in female flies, indicating potentially important sexual dimorphism in infection avoidance. Understanding how avoidance behaviours may vary is therefore important for our understanding of how disease will spread in natural populations ^4^, and more broadly how pathogens might evolve in response to variation in host infection avoidance strategies ^36,37^.

## Acknowledgements

We thank H. Cowan and H. Borthwick for preparing fly medium, K. Monteith and S. Lewis for assistance with the qPCR, the members of the Vale, Obbard and Ross labs for helpful suggestions. This work was supported by a strategic award from the Wellcome Trust for the Centre for Immunity, Infection and Evolution (grant reference no. 095831), and by a Society in Science - Branco Weiss fellowship, both awarded to P. Vale. P.Vale is also supported by a Chancellors Fellowship from the School of Biological Sciences, University of Edinburgh.

## Author contributions

PFV conceived the study. PFV and MDJ designed the experiment. MDJ and PFV carried out the experimental work. PFV analysed the data, wrote the manuscript and provided all research consumables.

## References

1. Fellous, S., Duncan, A. B., Quillery, E., Vale, P. F. & Kaltz, O. Genetic influence on disease spread following arrival of infected carriers. Ecol. Lett. 15, 186–192 (2012).

2. Vale, P. F. & Little, T. J. Measuring parasite fitness under genetic and thermal variation. Heredity 103, 102–109 (2009).

3. Susi, H., Barrès, B., Vale, P. F. & Laine, A.-L. Co-infection alters population dynamics of infectious disease. Nat. Commun. 6,(2015).

4. Barron, D., Gervasi, S., Pruitt, J. & Martin, L. Behavioral competence: how host behaviors can interact to influence parasite transmission risk. Curr. Opin. Behav. Sci. 6, 35–40 (2015).

5. Paull, S. H. et al. From superspreaders to disease hotspots: linking transmission across hosts and space. Front. Ecol. Environ. 10, 75–82 (2012).

6. Webster, C. L. et al. The Discovery, Distribution, and Evolution of Viruses Associated with Drosophila melanogaster. PLOS Biol 13, e1002210 (2015).

7. Curtis, V. A. Infection-avoidance behaviour in humans and other animals. Trends Immunol. 35, 457–464 (2014).

8. Parker, B. J., Barribeau, S. M., Laughton, A. M., de Roode, J. C. & Gerardo, N. M. Non-immunological defense in an evolutionary framework. Trends Ecol. Evol. 26, 242–248(2011).

9. Moore, J. An overview of parasite-induced behavioral alterations - and some lessons from bats. J. Exp. Biol. 216, 11–17 (2013).

10. Wolinska, J. & King, K. C. Environment can alter selection in host-parasite interactions. Trends Parasitol. 25, 236–244 (2009).

11. Vale, P. F., Salvaudon, L., Kaltz, O. & Fellous, S. The role of the environment in the evolutionary ecology of host parasite interactions. Infect. Genet. Evol. 8, 302–305 (2008).

12. Adamo, S. A. Comparative psychoneuroimmunology: evidence from the insects. Behav. Cogn. Neurosci. Rev. 5, 128–140 (2006).

13. Adelman, J. S. & Martin, L. B. Vertebrate sickness behaviors: Adaptive and integrated neuroendocrine immune responses. Integr. Comp. Biol. 49, 202–214(2009).

14. Parker, B. J., Elderd, B. D. & Dwyer, G. Host behaviour and exposure risk in an insect-pathogen interaction. J. Anim. Ecol. 79, 863–870 (2010).

15. Neyen, C., Bretscher, A. J., Binggeli, O. & Lemaitre, B. Methods to study Drosophila immunity. Methods San Diego Calif 68, 116–128 (2014).

16. Dubnau, J. Behavioral Genetics of the Fly (Drosophila Melanogaster). (Cambridge University Press, 2014).

17. Sokolowski, M. B. Drosophila: Genetics meets behaviour. Nat. Rev. Genet. 2, 879–890 (2001).

18. Dostert, C. et al. The Jak-STAT signaling pathway is required but not sufficient for the antiviral response of drosophila. Nat. Immunol. 6, 946–953 (2005).

19. Ferreira, Á. G. et al. The Toll-Dorsal Pathway Is Required for Resistance to Viral Oral Infection in Drosophila. PLoS Pathog. 10,(2014).

20. Huszar, T. & Imler, J. in Advances in Virus Research Volume 72, 227–265 (Academic Press, 2008).

21. Kapun, M., Nolte, V., Flatt, T. & Schlötterer, C. Host Range and Specificity of the Drosophila C Virus. PLoS ONE 5, e12421 (2010).

22. Arnold, P. A., Johnson, K. N. & White, C. R. Physiological and metabolic consequences of viral infection in Drosophila melanogaster. J. Exp. Biol. 216, 3350–3357(2013).

23. Chtarbanova, S. et al. Drosophila C virus systemic infection leads to intestinal obstruction. J. Virol. (2014). doi:10.1128/JVI.02320-14

24. Vale, P. F. & Jardine, M. D. Sex-specific behavioural symptoms of viral gut infection and Wolbachia in Drosophila melanogaster. J. Insect Physiol. 82, 28–32 (2015).

25. Livak, K. J. & Schmittgen, T. D. Analysis of relative gene expression data using real-time quantitative PCR and the 2(-Delta Delta C(T)) Method. Methods San Diego Calif 25, 402–408 (2001).

26. JMP. (SAS Institute Inc.).

27. Barber, I. & Dingemanse, N. J. Parasitism and the evolutionary ecology of animal personality. Philos. Trans. R. Soc. Lond. B Biol. Sci. 365, 4077–4088 (2010).

28. Ayres, J. S. & Schneider, D. S. The Role of Anorexia in Resistance and Tolerance to Infections in Drosophila. PLoSBiol 7, e1000150 (2009).

29. Behringer, D. C., Butler, M. J. & Shields, J. D. Ecology: Avoidance of disease by social lobsters. Nature 441, 421–421 (2006).

30. Babin, A. et al. Fruit flies learn to avoid odours associated with virulent infection. Biol. Lett. 10, 20140048 (2014).

31. Meisel, J. D. & Kim, D. H. Behavioral avoidance of pathogenic bacteria by Caenorhabditis elegans. Trends Immunol. 35, 465–470 (2014).

32. Hutchings, M.., Knowler, K.., McAnulty, R. & McEwan, J.. Genetically resistant sheep avoid parasites to a greater extent than do susceptible sheep. Proc. R. Soc. B Biol. Sci. 274, 1839–1844 (2007).

33. Curtis, V., de Barra, M. & Aunger, R. Disgust as an adaptive system for disease avoidance behaviour. Philos. Trans. R. Soc. Lond. B. Biol. Sci. 366, 389–401 (2011).

34. Kacsoh, B. Z., Lynch, Z. R., Mortimer, N. T. & Schlenke, T. A. Fruit Flies Medicate Offspring After Seeing Parasites. Science 339, 947–950 (2013).

35. Longdon, B., Cao, C., Martinez, J. & Jiggins, F. M. Previous Exposure to an RNA Virus Does Not Protect against Subsequent Infection in Drosophila melanogaster. PLoSONE 8, e73833 (2013).

36. Boots, M. & Bowers, R. G. Three mechanisms of host resistance to microparasites-avoidance, recovery and tolerance-show different evolutionary dynamics. J. Theor. Biol. 201, 13–23 (1999).

37. McLeod, D. V. & Day, T. Pathogen evolution under host avoidance plasticity. Proc. Biol. Sci. 282,(2015).

